# ABA homeostasis and long-distance translocation is redundantly regulated by ABCG ABA importers

**DOI:** 10.1101/2021.05.12.443788

**Authors:** Yuqin Zhang, Himabindu Vasuki, Jie Liu, Hamutal Bar, Shani Lazary, Aiman Egbaria, Dagmar Ripper, Laurence Charrier, Asaph Aharoni, Laura Ragni, Lucia Strader, Nir Sade, Roy Weinstain, Markus Geisler, Eilon Shani

## Abstract

The effects of abscisic acid (ABA) on plant growth, development and response to the environment depend on local ABA concentrations. Here, we exploited a genome-scale amiRNA screen, targeting the *Arabidopsis* transportome, to show that ABA homeostasis is regulated by two previously unknown ABA transporters. ABCG17 and ABCG18 are localized to the plasma membranes of leaf mesophyll and stem cortex cells to redundantly promote ABA import, leading to conjugated inactive ABA sinks, thus restricting stomatal closure. *ABCG17* and *ABCG18* double knockdown revealed that the transporters encoded by these genes not only limit stomatal aperture size, conductance and transpiration while increasing water-use efficiency but also control ABA translocation from the shoot to the root to regulate lateral root emergence. The proposed ABCG17- and ABCG18-dependent ABA glucosyl ester shoot sink mechanism is restrained under abiotic stress conditions to further activate the ABA responses.

## Introduction

Abscisic acid (ABA) is a plant hormone that regulates growth and responses to the changing environment. For example, seed dormancy, germination, drought tolerance, stomatal closure and lateral root emergence are modulated by this important hormone under normal conditions and in response to stimuli ^1, 2, 3, 4, 5, 6, 7^. The ABA response is regulated at multiple steps: biosynthesis, transport, catabolism, perception, and signal transduction. ABA travels long distances throughout the plant, a characteristic that was shown about 50 years ago to affect stomatal conductance and responses to drought ^8, 9, 10, 11^. For decades, the concept stated in textbooks was that in response to drought, root-derived ABA travels to shoot guard cells via the xylem sap to prevent water loss ^12, 13^. Indeed, several studies support the idea that root-derived ABA synthesis is required for ABA-induced stomatal closure to elicit drought tolerance ^14, 15, 16^. Recent studies, however, provide evidence that stomatal closure is triggered by shoot-specific ABA synthesis ^17, 18, 19, 20, 21, 22^.

Multiple lines of evidence support the hypothesis that ABA transport is necessary for proper ABA responses ^23, 24, 25^. First, ABA, being a weak acid, exists primarily in its charged and membrane-impermeable form, suggesting the need for transporter-facilitated movement across membranes ^26, 27, 28, 29, 30^. Second, using deuterium-labeled ABA and reciprocal grafting between wild-type and ABA-biosynthetic mutant tomatoes, it was found that foliage-derived ABA promotes root growth but inhibits the development of lateral roots ^31^. This highlights the physiological and morphological importance of long-distance ABA transport beyond the control of stomata. Third, phloem-specific ABA synthesis complements ABA activity in stomatal aperture, indicating that ABA movement within the leaf is important ^32, 33, 34^. Fourth, ABA transport is required for processes such as germination ^24, 35, 36^.

In recent years, several ABA transporters have been identified and characterized ^15, 24, 37, 38^. ABCG25 and ABCG31 exporters, members of the ATP-BINDING CASSETTE-G (ABCG), are involved in ABA movement out of the vasculature, endosperm, and guard cells. ABCG40 and ABCG30 are ABA importers, that transport ABA into the guard cells and into the embryo ^24, 37, 39, 40, 41^. MtABCG20 was recently characterized as an ABA importer that is involved in germination and lateral-root formation in *Medicago truncatula* ^42^. Additional transporters include DTX50, a transporter of the multidrug and toxic compound extrusion (MATE) transporter family, and NITRATE TRANSPORT1/PEPTIDE TRANSPORTER FAMILY (NPF) 4.6 (also known as NRT1.2 and AIT1) ^15, 24, 26, 34, 43, 44^.

In addition to transport, ABA concentrations are tightly controlled by ABA metabolism ^36, 45^. ABA is conjugated with glucose by UDP-glucosyltransferases (ABA-glucosyltransferase) to catalyze formation of the stored form of ABA, ABA-glucosyl ester (ABA-GE) ^46^. Previous studies showed that there are 26 subfamilies of UDP-glucosyltransferases (UGTs) in *Arabidopsis*. Seven UGTs, which belong to different groups of UDPs subfamily 1 (encoded by *UGT84B1, UGT75B1, UGT84B2, UGT71B6, UGT75B2, UGT73B1*, and *UGP71C5*) have ABA to ABA-GE catalysis activity ^47, 48^. However, our understating on the spatio-temporal aspects of ABA homeostasis and on the mechanisms underlying ABA translocation to control ABA responses under normal and stress conditions are limited.

Here we exploited a genome-scale amiRNA screen targeting the *Arabidopsis* transportome to identify novel ABCG ABA transporters. ABCG17 and ABCG18 are localized to the plasma membrane and import ABA. *ABCG17*- and *ABCG18*-knockdown lines displayed enhanced ABA accumulation, induced ABA response, and reduced stomatal aperture size, conductance, and transpiration with increased water-use efficiency compared to WT. In addition, seedlings with double-loss-of-function of *ABCG17* and *ABCG18* showed enhanced long-distance ABA translocation to the root, which led to lateral root outgrowth inhibition. ABCG17 and ABCG18 are primarily expressed in the shoot mesophyll and cortex cells, where they drive ABA import, allowing ABA-GE formation. In turn, these ABA-GE sinks prevent active ABA accumulation in guard cells and limit long-distance travel of ABA to lateral root formation sites. Under abiotic stress conditions, *ABCG17* and *ABCG18* are transcriptionally repressed, promoting active ABA movement, accumulation, and response. The transport mechanism mediated by ABCG17 and ABCG18 allows plants to maintain ABA homeostasis under normal growth conditions.

## Results

### Transportome-scale amiRNA screen indicates involvement of class A ABCGs in ABA mediated activity

Over 75% of the genes in the *Arabidopsis* genome belong to gene families, complicating functional analyses ^49^. In order to partially overcome functional redundancy and reveal genetic factors involved in ABA homeostasis and translocation, we utilized a transportome-specific amiRNAs collection of 3,000 lines targeting multiple transporters from the same family ^50^. The transporter amiRNA sub-library is part of the broad PHANTOM amiRNA tool ^49^. The seedlings in the amiRNA collection were screened for defects in shoot growth and photosynthesis-related parameters. The screen identified *amiRNA-1228* line as an interesting candidate; this line had slight but significant shoot growth inhibition (Fig. 1a, Supplementary Fig. 1a). In addition, the *amiRNA-1228* line showed significant reduction in photosystem II quantum yield of light-adapted sample at steady state (Fig. 1b), indicating limited activity of photosystem II ^51^. We therefore hypothesized that the genes targeted by *amiRNA-1228* might be related to ABA activity and homeostasis.

**Fig. 1.**
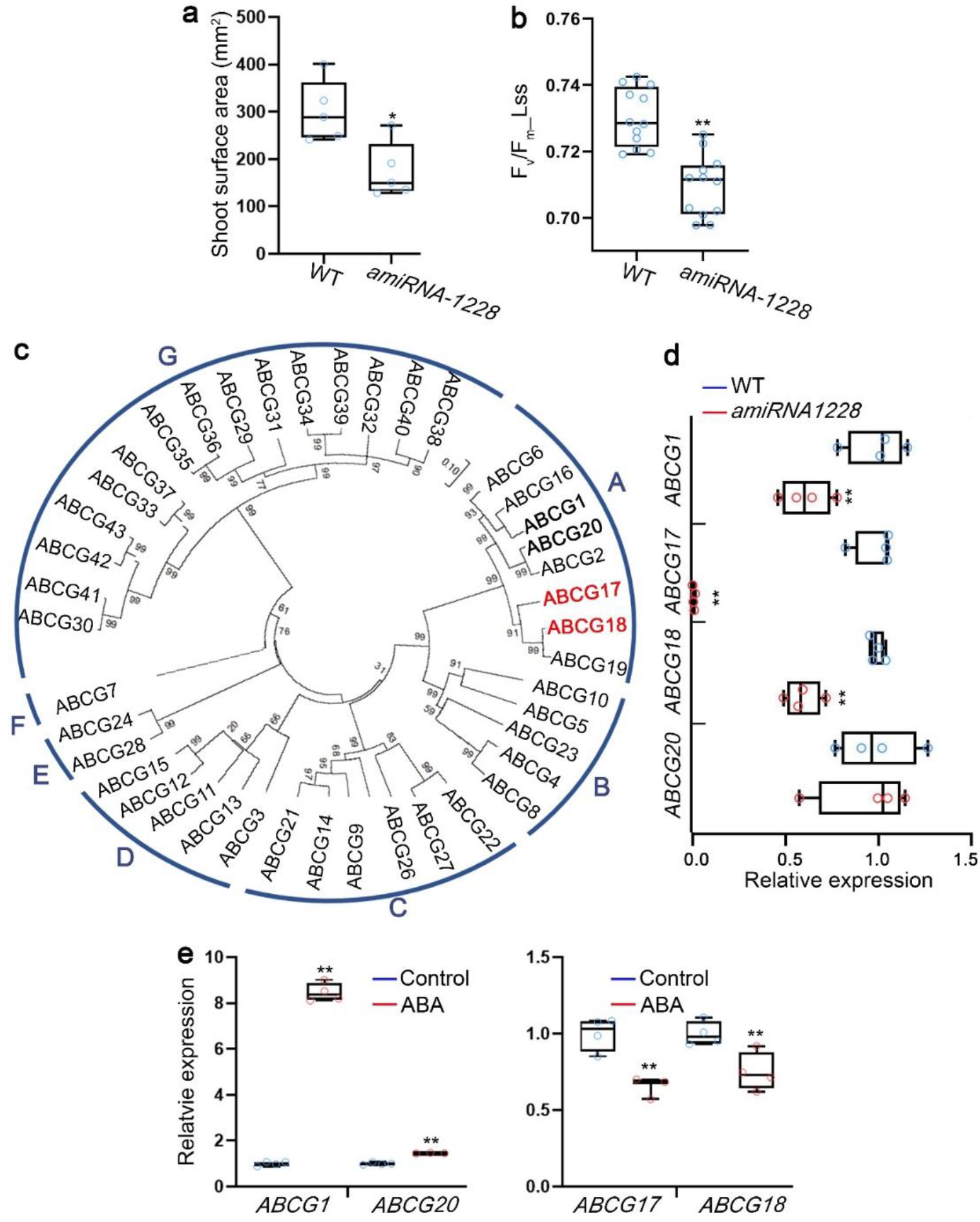
Reduced expression of several class A ABCG members results in moderate shoot growth and decreased photosynthesis rate. **a,** Shoot surface area of 18-day-old WT and *amiRNA-1228*. n = 5, *P < 0.05, Student’s t-test. **b,** Fv/Fm_Lss, indicating for photosystem II quantum yield of light-adapted sample at steady state, measured for 18-day-old WT and *amiRNA-1228.* Shown are averages (±SE), n = 12, ** P < 0.01, Student’s t-test. **c,** Phylogenetic tree of *Arabidopsis* ABCG family based on amino acid sequences. Red and bold fonts indicate proteins coded by putative *amiRNA-1228* target genes. **d,** Relative expression of the indicated *ABCG amiRNA-1228* targeted genes in 12-day-old WT and *amiRNA-1228* seedlings, quantified by qRT-PCR. n = 4, **P < 0.01, Student’s t-test. **e,** Relative expression, quantified by qRT-PCR, of the indicated *ABCG amiRNA-1228* targeted genes in response to ABA treatment (5 μM ABA for 3 hours) in 7-day-old seedlings. n = 4, **P < 0.01, Student’s t-test.

*amiRNA-1228* putatively targets four class A ABCG subfamily members with different extents of complementarity in the potential recognition sites (Supplementary Fig. 1b). ABCG17 and ABCG18 are grouped in one branch, ABCG1 and ABCG20 are classified into another, of the *Arabidopsis* ABCG family phylogenetic tree (Fig. 1c). To test if these genes are indeed targeted by *amiRNA-1228*, we examined their expression levels by qRT-PCR in *amiRNA-1228* plants compared to levels in wild-type (WT) plants (Fig. 1d). *ABCG1, ABCG1*7, and *ABCG18* were significantly down-regulated in *amiRNA-1228*, but there was no difference in levels of *ABCG20* (Fig. 1d). Next, we tested the transcriptional responses of these genes to ABA treatment. *ABCG1* and *ABCG20* were induced by 5 μM ABA treatment for 3 hours, whereas *ABCG17* and *ABCG18* were repressed (Fig. 1e). ABCG1 and ABCG16 were previously shown to regulate pollen and reproductive development ^52, 53^ and suberin formation in roots ^54^. ABCG20, ABCG2, and ABCG6 regulate suberin barriers in roots and seed coats ^53^. The functions of ABCG17 and ABCG18, located on separate phylogenetic branch (Fig. 1c), have not been characterized. Altogether, our results indicate that *ABCG17* and *ABCG18* genes are interesting candidates in the context of ABA-mediated activity.

### *ABCG17* and *ABCG18* are redundantly required for ABA response and stomatal closure

To determine whether *ABCG17* and *ABCG18* contribute to the *amiRNA-1228* phenotype and to dissect their activity, we obtained T-DNA insertion lines for both genes (Fig. 2a). Since *ABCG17* and *ABCG18* are genetically linked on chromosome 3, we could not generate double mutants by crossing T-DNA-insertion lines. Therefore, two independent amiRNA double-knockdown lines for *ABCG17* and *ABCG18* were generated. The first was an amiRNA targeting both *ABCG17* and *ABCG18* (*mir17,18*). The second was an amiRNA targeting *ABCG17* (*mir17*) transformed into the *abcg18-1* T-DNA-insertion background (*mir17,g18-1*) (Fig. 2a). We examined the expression levels of *ABCG17* and *ABCG18* in the single- and double-knockdown lines compared to WT. Both *ABCG17* and *ABCG18* were significantly downregulated in the respective backgrounds (Fig. 2b-c). Next, we monitored the shoot phenotypes of *abcg17* and *abcg18* single-mutant and double-knockdown lines grown in soil under normal growth conditions. None of the single mutants showed shoot-growth impairment, but both double-knockdown lines, *mir17,18* and *mir17,g18-1*, displayed significant reductions in shoot surface area compared to WT and to the single mutants (Fig. 2d-e). Similarly, 50-day-old double-knockdown plants had shorter inflorescence stems compared to WT and single mutants (Fig. 2f-g). The results are in line with the *amiRNA-1228* phenotype and indicate that *ABCG17* and *ABCG18* are redundantly required for plant shoot growth.

**Fig. 2.**
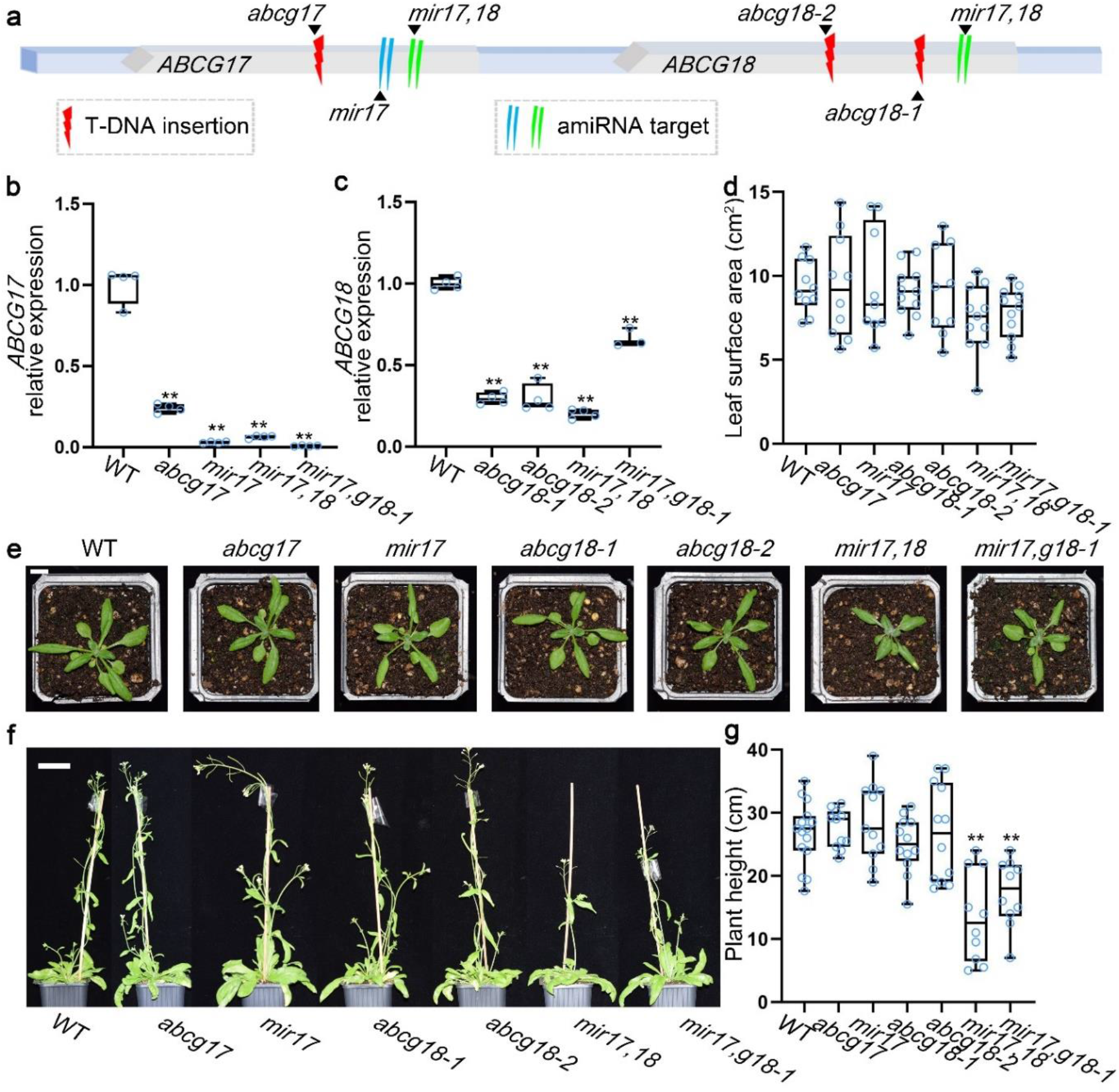
*ABCG17* and *ABCG18* redundantly regulate plant growth. **a,** Illustration of the *ABCG17* and *ABCG18* genome region. T-DNA insertions and amiRNA knockdown sites are indicated. **b-c,** Relative expression levels of *ABCG17* (b) and *ABCG18* (c) in the indicated genotypes. *mir17,18* is the *abcg17, abcg18* double-knockdown amiRNA line; *mir17,g18-1* is *mir17 (amiRNA-ABCBG17)* transformed into the background of *abcg18-1* T-DNA insertion line. n = 4, **P < 0.01, Student’s t-test. **d,** Leaf surface area of the indicated lines. n ≥ 9 plants, **P < 0.01, Student’s t-test. **e,** Shoot phenotypes of 25-day-old plants grown in soil under normal conditions. Scale bar = 1 cm. **f,** Phenotypes of 50-day-old plants grown in soil under normal conditions. Scale bar = 2 cm. **g,** Quantification of inflorescence stem height of the indicated genotypes. Shown are averages (±SE), n ≥ 10 plants, **P < 0.01, Student’s t-test.

Since *amiRNA-1228* showed chlorophyll fluorescence related phenotypes, we tested whether *ABCG17* and *ABCG18* contribute to this activity. Single *abcg17* and *abcg18* mutants did not differ from WT plants in stomatal aperture size. However, both *mir17,18* and *mir17,g18-1* had significantly smaller stomatal apertures than WT and single-mutant plants (Fig. 3a). Similarly, the double-knockdown lines had significantly reduced stomatal conductance compared to WT and single mutants (Fig. 3b). Next we tested whether gas exchange (i.e. transpiration and photosynthesis) rates were affected in the mutant lines. In line with stomatal conductance and aperture defects, the double-knockdown lines had significantly lower transpiration rates than WT and single mutants (Fig. 3c). However, photosynthesis rates were not changed within experimental error in both single mutants and the double-mutant *mir17,18* line compared to WT plants (Fig. 3d). In agreement with this finding, the double-mutant line, *mir17,18* showed an increase in instantaneous water use efficiency under well-watered conditions (Fig. 3e). Analysis of water-loss rates from excised leaves of single mutants and *mir17,18* showed delayed water-loss rates compared to WT when exposed to air (Fig. 3f, Supplementary Fig. 2). Evaluation of leaf temperature utilizing thermal imaging showed higher leaf temperature in both the double-knockdown lines compared to WT and single mutants (Fig. 3g-h). These data support the hypothesis that ABCG17 and ABCG18 are required for stomata activity under normal conditions.

**Fig. 3.**
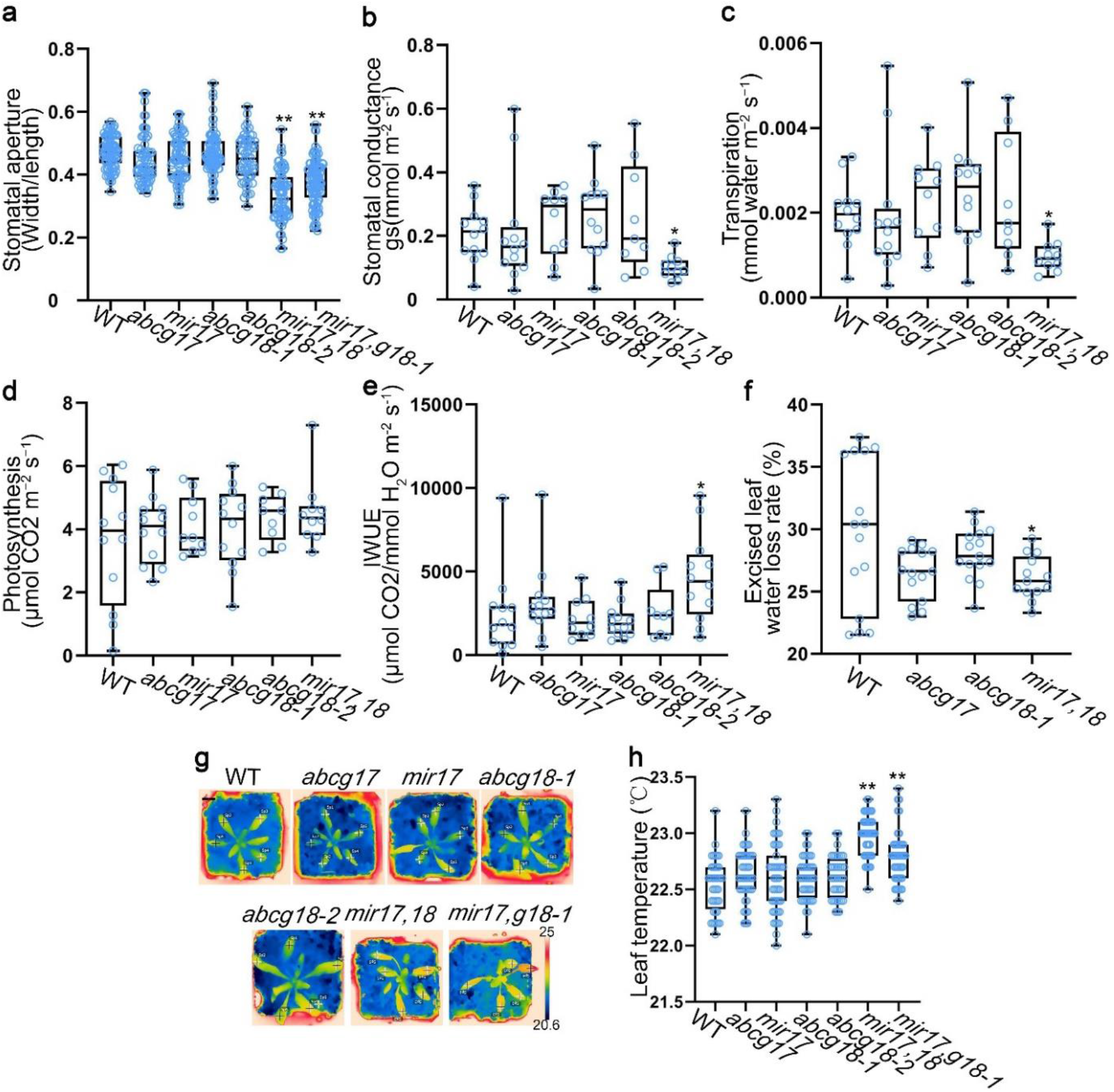
Stomatal aperture, conductance, and transpiration is regulated by *ABCG17* and *ABCG18*. **a,** Stomatal aperture measurements of 25-day-old plants of the indicated genotypes. *mir17,18* is *abcg17, abcg18* double-knockdown amiRNA line, *mir17,g18-1* is *mir17* (*amiRNA-ABCBG17*) transformed into the background of *abcg18-1* T-DNA line. Shown are averages (±SE), n ⩾ 45, **P < 0.01, Student’s t-test. **b,** Plant stomatal conductance measurements of 25-day-old plants of the indicated genotypes. n = 5 plants, *P < 0.05, Student’s t-test. **c,** Leaf transpiration rates of the 25-day-old plants of the indicated genotypes. n = 5 plants, *P < 0.05, Student’s t-test. **d,** Leaf photosynthetic properties of 25-day-old plants of the indicated genotypes. n = 5 plants. Results were not significant at P > 0.05 by Student’s t-test. **e,** Instantaneous water use efficiency (IWUE) of plants of the indicated genotypes under normal conditions. n = 5 plants, *P < 0.05, Student’s t-test. **f,** Water-loss rates of leaves excised from 30-day-old plants of the indicated genotypes (90 minutes, room temperature air). n ⩾ 13. *P < 0.05, Student’s t-test. **g,** Thermal images of 25-day-old plants of the indicated genotypes grown on soil under normal conditions. “+” indicates leaf temperature quantification sites. Scale bar = 1 cm. **h,** Leaf temperatures of plants of the indicated genotypes measured at sites indicated in panel (g). n ⩾ 40, *P < 0.01, Student’s t-test.

To further characterize ABCG17 and ABCG18 activity, we generated transgenic plants overexpressing (*35S* promoter) each of the two genes. Two independent lines for each gene showed 50-100-fold change in transcriptional activation (Supplementary Fig. 3a-b). *ABCG17-* or *ABCG18-*overexpressing lines showed no differences in shoot surface area from that of WT plants, but *ABCG18* overexpression resulted in a slight plant height increase (Supplementary Fig. 3c-f). *ABCG17* or *ABCG18* overexpression resulted in reduced stomatal aperture (Supplementary Fig. 4a-b), elevated leaf temperature (Supplementary Fig. 4c-d), and induced ABA response, reflected by the induction of *pRAB18:GFP* ^55^ and *pMAPKKK18:LUC ^56^* ABA reporters (Supplementary Fig. 4e-f) compared to WT plants. Together, these results suggest that ABCG17 and ABCG18 redundantly regulate stomatal conductance and aperture.

### ABCG17 and ABCG18 are plasma membrane localized ABA importers

To determine the subcellular localizations of ABCG17 and ABCG18, we cloned the coding sequences of *ABCG17* and *ABCG18* and constructed vectors for expression of these proteins with N-terminal YFP tags under the control of the *35S* promoter. Confocal microscopy of *p35S:YFP-ABCG17* and *p35S:YFP-ABCG18* transgenic lines revealed that ABCG17 and ABCG18 are localized to the plasma membrane (Fig. 4a).

**Fig. 4.**
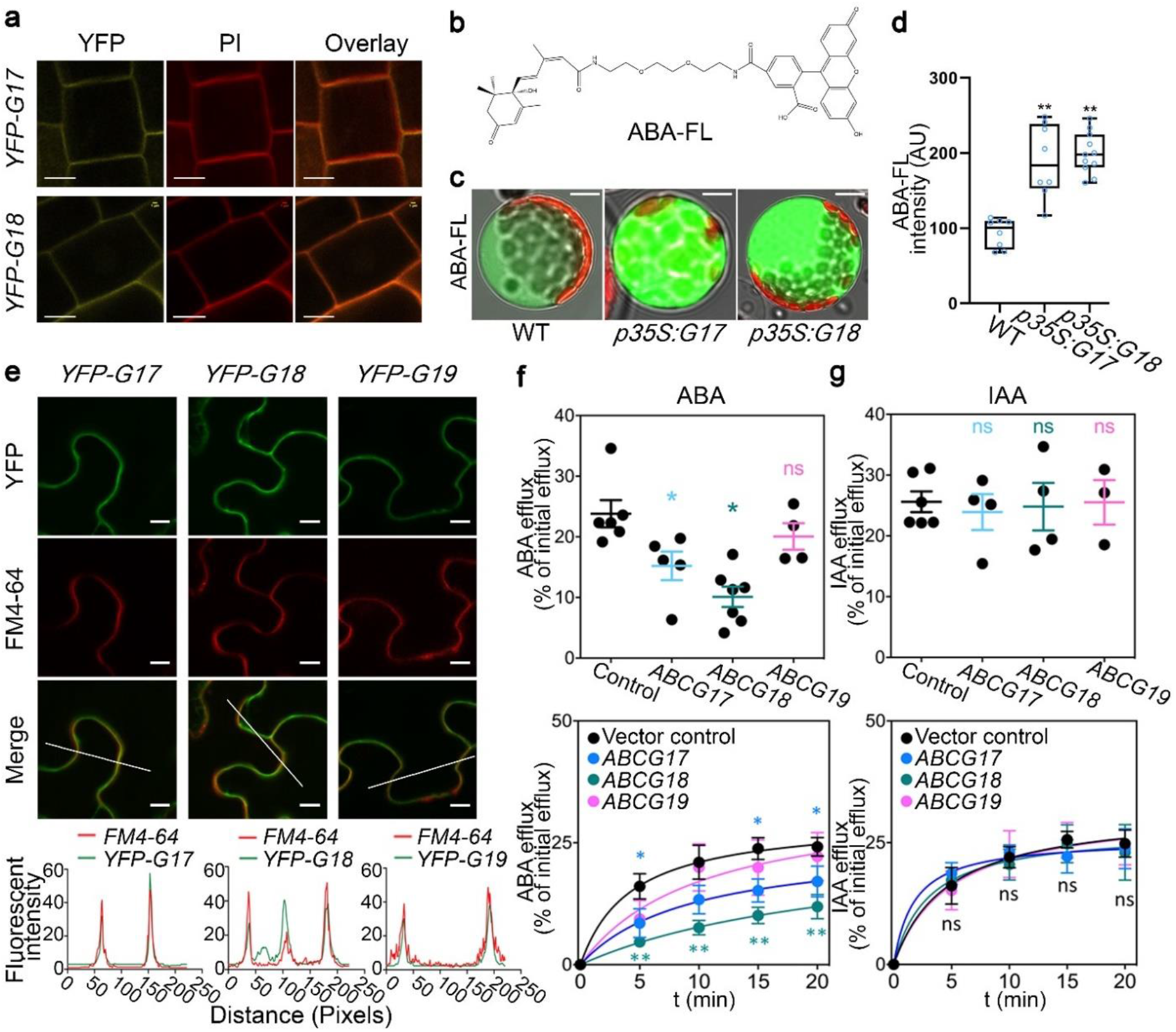
ABCG17 and ABCG18 are ABA importers localized to the plasma membrane. **a,** ABCG17 and ABCG18 subcellular localization shown by *p35S:YFP-ABCG17* (*YFP-G17*) and *p35S:YFP-ABCG18* (*YFP-G18*) stable transgenic *Arabidopsis* lines. Fluorescence was imaged in the root meristem epidermis layer. Yellow indicates YFP-ABCG17 or YFP-ABCG18 fluorescence, red indicates propidium iodide (PI). Scale bar = 5 μm. **b,** Molecular structure of ABA-FL, a fluorescein moiety is linked to the ABA carboxylic group. **c,** ABA-FL fluorescent signal in *Arabidopsis* protoplasts overexpressing *ABCG17* or *ABCG18*. *p35S:G17* and *p35S:G18* indicate *p35S:ABCG17* and *p35S:ABCG18*. Protoplasts were treated with 5 μM ABA-FL for 15 min. Scale bar = 10 μm. **d,** Quantification of ABA-FL fluorescence intensity in protoplasts overexpressing *ABCG17* or *ABCG18*. n ≥ 8 protoplasts, **P < 0.01, Student’s t-test. **e,** Confocal imaging of tobacco leaves transfected with *p35S:YFP-ABCG17* (*YFP-G17*), *p35S:YFP-ABCG18* (*YFP-G18*) or *p35S:YFP-ABCG19* (*YFP-G19*), stained with FM4-64 plasma membrane marker. Scale bar = 10 μm. Bottom graphs are quantified fluorescence intensities of pixels along the white line within the images. X-axis represents distance along the line. **f,** [^14^C]ABA efflux from tobacco protoplasts as percentage of initial efflux. Top graph is 15 min [^14^C]ABA, n ≥ 4, *P < 0.05, Welch’s t-test. **g,** [^3^H]IAA efflux from tobacco protoplasts as percentage of initial efflux. Top graph is 15 min [^3^H]IAA, n ≥ 4, ns indicates not significant.

To examine whether ABCG17 and ABCG18 facilitate ABA transport, we synthesized fluorescently labeled ABA molecules by adding a fluorescein-tag to the carboxylic group (Fig. 4b). Unlike the fluorescently labeled gibberellin molecules we previously reported ^57^, the fluorescent ABA (ABA-FL) was only slightly bioactive in root growth assays, suggesting low binding affinity for the PYR/PYL/RCAR ABA receptor (Supplementary Fig. 5). We used the new ABA-FL molecule to test ABCG17 and ABCG18 transport activity. Isolated protoplasts from leaf mesophylls of transgenic *Arabidopsis* plants overexpressing *ABCG17* or *ABCG18*, and WT as a control were treated with 5 μM ABA-FL for 12 hours and imaged. Protoplasts overexpressing *ABCG17* or *ABCG18* had enhanced ABA-FL fluorescent signal compared to WT protoplasts (Fig. 4c-d), suggesting that both *ABCG17* and *ABCG18* promote ABA import.

To validate ABCG17 and ABCG18 ABA transport activity, a biochemical transport assay with radiolabeled ABA ([^14^C]ABA) in tobacco plants that overexpress each of these two transporters was carried out. *ABCG17* or *ABCG18* overexpression in tobacco leaves showed plasma membrane localization (Fig. 4e) and resulted in significant reduction in measured protoplast export activity compared to the control (Fig. 4f). Reduced net export is in line with an import activity for both transporters that would function here as re-importers of [^14^C]ABA. In order to determine the specificity of ABCG17 and ABCG18 in ABA transport activity, we tested the effects of overexpressing *ABCG19*, the closest gene in the cluster (Fig. 1c). Notably, overexpression of *ABCG19* did not result in significant ABA transport activity (Fig. 4f). We also evaluated transport of a radioactively labeled auxin ([^3^H]IAA), like ABA, an organic acid and a known substrate of members of the ABCB subclass. *ABCG17* or *ABCG18* overexpressing lines did not reveal altered transport of auxin compared to the control (Fig. 4g). These experiments indicate that ABCG17 and ABCG18 are plasma membrane localized ABA importers.

### ABCG17 and ABCG18 limit long-distance ABA transport to regulate lateral root development

In addition to the stomatal conductance and aperture size phenotypes, double knockdown of *ABCG17* and *ABCG18* resulted in reduced lateral root numbers compared to WT and single mutants (Fig. 5a). More specifically, *mir17,18* showed normal lateral root initiation (Supplementary Fig. 6) but reduced lateral root primordia outgrowth compared to WT (Fig. 5b). This result is intriguing since expression of *ABCG17* and *ABCG18* is largely restricted to the shoot: qPCR measurements, luciferase reporter lines (*pABCG17:LUC* and *pABCG18:LUC*), GUS reporter lines (*pABCG17:GUS* and *pABCG18:GUS*), and YFP reporter lines (*pABCG17:NLS-YFP* and *pABCG18:NLS-YFP*) showed strong signals in shoots (Fig. 5c-e). A very weak signal was found in lateral root emerging primordia using the *pABCG17:NLS-YFP pABCG18:NLS-YFP* lines, only when applying maximum laser power and using a high sensitivity gallium arsenide phosphide (GaAsP) detector (Fig. 5f, Supplementary Fig. 7a-b). ABA regulates key stages of lateral root post-emergence development ^58^. Specifically, ABA signaling plays crucial role in salt stress-induced lateral root suppression ^5, 59^ and lateral root growth recovery ^60^, associated with an increase in *miR165a* and a reduction in *PHB* (*PHABULOSA*) levels ^61^.

**Fig. 5.**
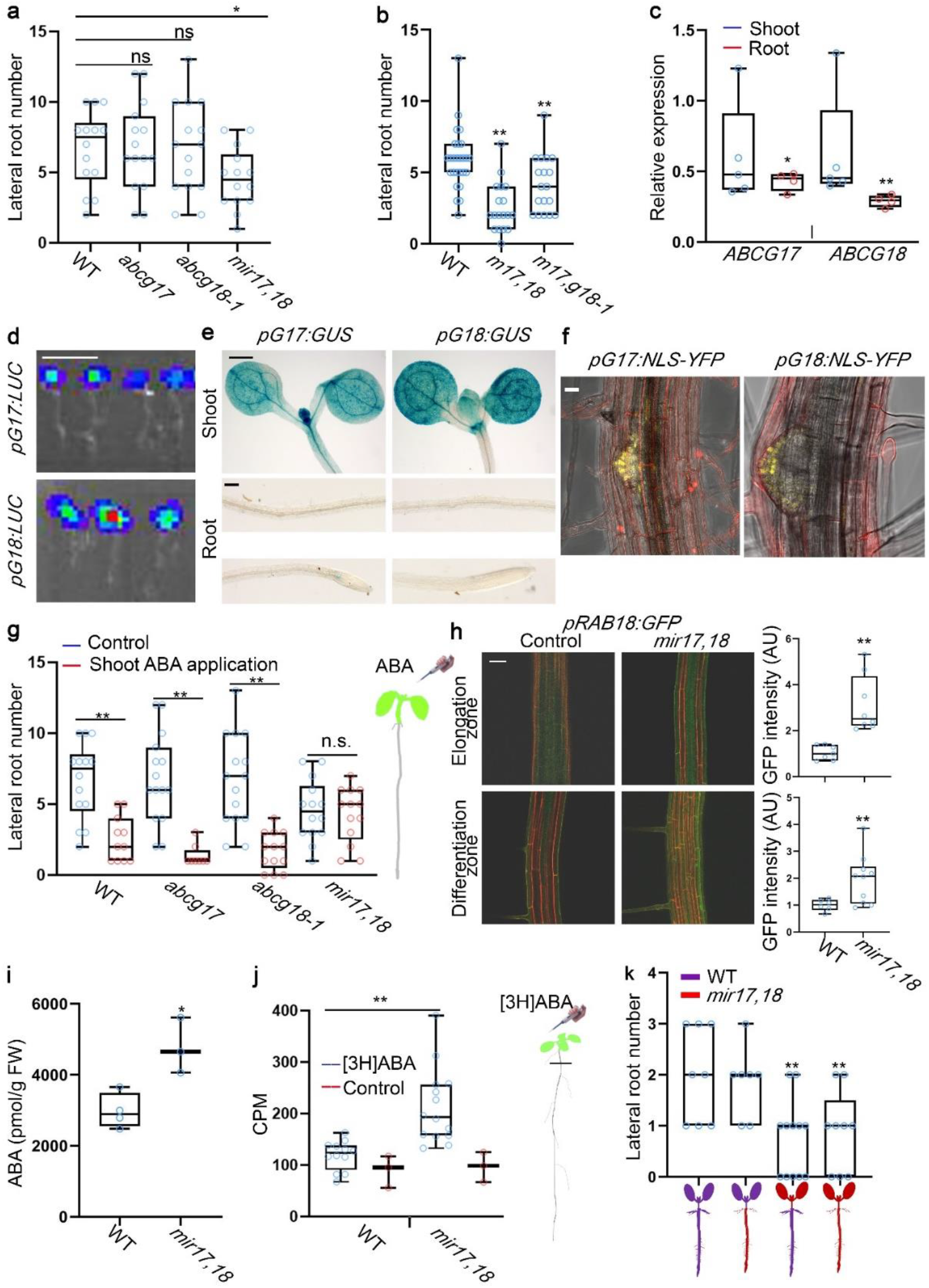
ABCG17 and ABCG18 are required for long-distance ABA transport and regulation of lateral root formation. **a,** Lateral root numbers of 10-day-old seedlings for indicated genotypes. *mir17,18* is the *abcg17, abcg18* double-knockdown amiRNA line. n ≥ 14, *P < 0.05, Student’s t-test. ns indicates for not significant. **b,** Lateral root numbers of 10-day-old plants for indicated genotypes. n ≥ 18, **P < 0.01, Student’s t-test. **c,** Relative expression of *ABCG17* and *ABCG18* in shoots and roots, quantified by qRT-PCR. n ≥ 4, **P < 0.01, *P < 0.05, Student’s t-test. **d,** Bioluminescence signal driven by *pABCG17:LUC* (*pG17:LUC*) and *pABCG18:LUC* lines (*pG18:LUC*). Scale bar = 1 cm. **e,** GUS staining for *pABCG17:GUS* (*pG17:GUS*) and *pABCG18:GUS* (*pG18:GUS)* in shoot and root of 5-day-old seedlings. Scale bar = 0.5 mm for shoots; scale bar = 0.1 mm for roots. **f,** Representative images of *pABCG17:NLS-YFP* (*pG17:NLS-YFP*) and *pABCG18:NLS-YFP* (*pG18:NLS-YFP*) lateral root emergence sites (YFP in yellow). Scale bar = 20 μm. **g,** Lateral root numbers for the indicated genotypes. Shoots of 4-day-old plants were treated with 5 μM ABA for 4 days. n ≥ 13, **P < 0.01, *P < 0.05, Student’s t-test. n.s. indicates not significant. To the right is an illustration of shoot-specific ABA application. **h,** *pRAB18:GFP* signal in the background of *miR17,18* and control roots (left) and the quantification of respective GFP signal intensity (right). Scale bar = 20 μm. **P < 0.01, Student’s t-test. **i,** Quantification of ABA amounts in roots of 12-day-old plants of the indicated genotypes. FW: fresh weight. n ≥ 3. *P < 0.05, Student’s t-test. **j,** [^3^H]ABA counts per minute (CPM) in roots only (roots were isolated as indicated by a black line in the illustration on the right). Controls were samples not treated with [^3^H]ABA. n ≥ 10, **P < 0.01, Student’s t-test. **k,** Lateral root number quantification of the indicated grafted lines, grafting performed on 6-day-old seedlings, lateral roots were scored 8 days after grafting. *mir17,18* is *abcg17, abcg18* double-knockdown amiRNA line. n ≥ 7, **P < 0.01, Student’s t-test compared to WT self-grafting plants.

In order to test if shoot-born ABA can affect lateral root development, we applied ABA specifically to shoots and quantified lateral root length and lateral root number. The results indicated that application of ABA to WT shoots inhibited lateral root elongation and lateral root development (Supplementary Fig. 8a-b). Moreover, ABA application to the shoots induced expression of the *pMAPKKK18:GUS* ABA reporter ^56^ in the root (Supplementary Fig. 8c). Shoot-specific ABA application inhibited lateral root emergence in the single mutants similarly to WT, but the *mir17,18* double knockdown mutant was insensitive to ABA application to shoots (Fig. 5g). We speculated that *mir17,18* accumulates high ABA levels in the root compared to WT under normal conditions.

To test this hypothesis, we analyzed the ABA reporter *pRAB18:GFP* in the background of *ABCG17* and *ABCG18* loss-of-function lines. We found enhanced *pRAB18:GFP* signal in *mir17,18* roots compared to control roots (Fig. 5h), supporting the hypothesis that *ABCG17* and *ABCG18* knockdown causes accumulation of abnormally high levels of ABA in the root. To confirm this hypothesis, we quantified ABA levels in roots. Indeed, the double-knockdown *mir17,18* plants accumulated significantly higher ABA amounts in the root compared to WT (Fig. 5i).

In order to further understand an apparent ABCG17 and ABCG18 activity in limiting ABA shoot-to-root translocation, we treated shoots with radioactively labeled ABA ([^3^H]ABA), and quantified counts in roots after 24 hours. The *mir17,18* roots accumulated higher levels of [^3^H]ABA than the WT roots (Fig. 5j), supporting the hypothesis that ABCG17 and ABCG18 activities restrict long-distance transport of ABA to the root. To further confirm this result, reciprocal WT and *mir17,18* grafting assays were carried out. The results showed that, within experimental error, WT scion/*mir17,18* rootstock grafts developed the same number of lateral roots as WT self-grafted plants. However, *mir17,18* scion/WT rootstock grafts developed significantly fewer lateral roots, with lateral root numbers similar to *mir17,18* self-grafted plants (Fig. 5k). These results indicate that ABCG17 and ABCG18 redundantly restrict shoot-to-root ABA translocation, mediating lateral root development under normal conditions.

### *ABCG17* and *ABCG18* are expressed in shoot mesophyll and cortex cells under non-abiotic stress conditions

The obtained results suggest that ABCG17 and ABCG18 limit ABA levels in guard cells and during lateral root emergence, through a process that takes place in the shoot. We therefore speculated that ABCG17 and ABCG18 redundantly regulate ABA homeostasis, possibly by allowing ABA storage. To test this hypothesis, we evaluated ABA and ABA-GE levels in shoots. The double-knockdown *mir17,18* plants accumulated higher ABA levels in the shoot than WT or single mutant plants (Fig. 6a), consistent with results showing that double-knockdown lines have reduced stomatal aperture size, conductance, reduced transpiration, and elevated leaf temperature (Fig. 3). Interestingly, the double-knockdown *mir17,18* shoots accumulated less ABA-GE than WT shoots (Fig. 6b), suggesting that ABCG17 and ABCG18 might promote ABA-GE production catalyzed by UDP-glucosyltransferases. In support of this hypothesis, the ABA response gene *ABI1* was significantly induced in *miR17,18* compared to WT plants (Fig. 6c). To better characterize *ABCG17* and *ABCG18* function in plant hormone metabolism, we quantified the metabolites hormone profile of additional ABAs, as well as gibberellins and auxins, two hormones largely antagonistic to ABA. The 37 metabolites profile included bioactive hormones, their precursors, catabolites and conjugated forms. The most profound affect was found for ABA and its inactive catabolites and conjugated forms (Fig. 6d, Supplementary Fig. S9). A weak and largely insignificant response was found for the gibberellin and auxin profile. The few metabolites that did change in the gibberellin and auxin profile were GA_24_, ICA, IET, IAA and IAA-Ala significantly higher in *abcg17* or *abcg18* single mutants and GA_44_ and IAA-Ala significantly higher in *mir17,18* (Fig. 6d, Supplementary Fig. S9). To identify the exact cell types that accumulate ABA due to transport facilitated by ABCG17 and ABCG18, we characterized the expression patterns of *ABCG17* and *ABCG18* in leaf and stem sections. *pABCG17:NLS-YFP* and *pABCG18:NLS-YFP* lines showed that *ABCG17* and *ABCG18* are primarily expressed in parenchyma shoot cells, which are the leaf mesophyll cells (Fig. 6e-f, Supplementary Fig. S10a-b). We did not detect expression in the leaf or stem veins or bundle sheath cells (Fig. 6e-f). These results imply that under normal conditions, ABCG17 and ABCG18 promote ABA uptake into shoot parenchyma cells, leading to increased ABA-GE levels. Thus, under non-stress conditions, low levels of free shoot-born ABA are available to guard cells and for long-distance movement to roots.

**Fig. 6.**
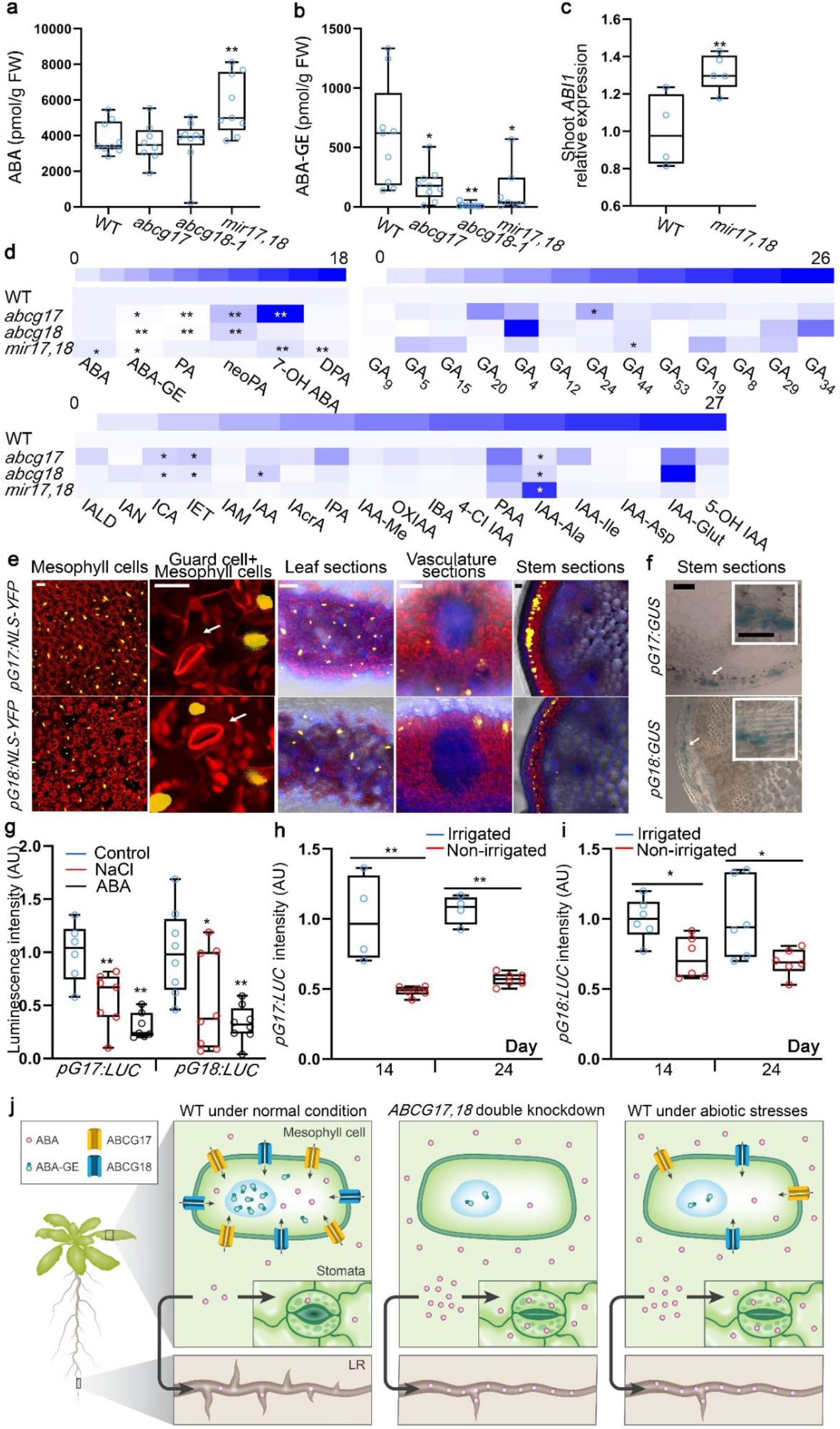
*ABCG17* and *ABCG18* are primarily expressed in mesophyll and cortex cells, allowing ABA-GE formation under normal conditions. **a-b,** Quantification of ABA (a) and ABA-GE (b) in shoots of 12-day-old plants of the indicated genotypes. FW: fresh weight. n ≥ 7. **P < 0.01, *P < 0.05, Student’s t-test compared to WT. **c,** Expression of *ABI1* in 12-day-old WT and *mir17,18* seedlings, quantified by qRT-PCR. n = 4, **P < 0.01, Student’s t-test. **d,** Metabolites profile heat-map for ABAs, gibberellins and auxins (bioactive, precursors, catabolites and conjugated forms) in shoots of 12-day-old plants of the indicated genotypes. Data is relative to WT. Full description of metabolites names and fold-change is shown in Supplementary S9. n ≥ 7. **P < 0.01, *P < 0.05, Student’s t-test. **e,** *pABCG17:NLS-YFP* (*pG17:NLS-YFP*) and *pABCG18:NLS-YFP* (*pG18:NLS-YFP*) signal (yellow) in leaves and stem. YFP signal is detected in mesophyll and cortex cells. Chlorophyll in red. Fluorescent brightener in blue. White arrows point to the guard cells. Scale bars = 20 μm. **f,** GUS staining for *pABCG17:GUS* (*pG17:GUS*) and *pABCG18:GUS* (*pG18:GUS*) in inflorescence stem of 30-day-old plants. Scale bar = 100 μm. Insets are magnifications of areas indicated by white boxes. Scale bar = 50 μm. **g,** Luminescence intensity of *pABCG17:LUC* (*pG17:LUC*) and *pABCG18:LUC* (*pG18:LUC*) with and without 5 μM ABA or 100 mM NaCl treatment. n ≥ 6, **P < 0.01, *P < 0.05, Student’s t-test, compared to control. **h-i,** Luminescence intensity of *pABCG17:LUC* (*pG17:LUC*) (h) and *pABCG18:LUC* (*pG18:LUC*) (i). Plants were irrigated for 15 days following by water withhold for 3 weeks. Control plants (irrigated) were watered throughout the experiment. n ≥ 4, **P < 0.01, *P < 0.05, Student’s t-test. **j,** Proposed model illustrating ABCG17 and ABCG18 function in regulating ABA homeostasis under normal and abiotic stress conditions. LR, lateral root.

Next, we wanted to elucidate how ABCG17 and ABCG18 function in ABA-mediated homeostasis under stress conditions, when high levels of ABA are needed. We therefore monitored the expression levels of *ABCG17* and *ABCG18* in response to abiotic stresses using *pABCG17:NLS-YFP* and *pABCG18:NLS-YFP* plants as well as *pABCG17:LUC* and *pABCG18:LUC* plants. *ABCG17* and *ABCG18* were transcriptionally repressed following ABA or NaCl treatments (Fig. 6g, Supplementary Fig. S11a-b), consistent with qRT-PCR experiments following ABA treatment (Fig. 1e). Similarly, long-period water-withhold experiments showed that *ABCG17* and *ABCG18* are repressed as intensity of luminescence due to the reporter expression was reduced in plants not irrigated for several days (7, 14 and 24 days) (Fig. 6h-i, Supplementary Fig. S11c). In summary, our experiments imply that ABCG17 and ABCG18 are necessary for ABA homeostasis through creation of ABA-GE sinks in mesophyll and cortex cells. These ABA-GE sinks limit ABA accessibility to guard cells and ABA availability for long-distance translocation to regulate lateral root formation. The ABA homeostasis shoot sink is released once the plant senses abiotic stress conditions.

## Discussion

Here we utilized a transportome-scale amiRNA screen to overcome functional redundancy to identify two previously unstudied ABCG transporters, *ABCG17* and *ABCG18*. Compared to WT and single *abcg17* and *abcg18* mutants, the double-knockdown lines had higher ABA content and reduced ABA-GE content in shoots. Since ABCG17 and ABCG18 are primarily expressed in the shoot cortex and mesophyll cells, localized to the plasma membrane, and promote ABA import, we hypothesized that the two proteins are redundantly required for ABA accumulation and storage in these cells (Fig. 6j). The ABCG17- and ABCG18-dependent ABA sink in mesophyll cells limits the hormone translocation and activity to guard cells and the lateral-root emergence sites, two distinct ABA-mediated processes. The two redundant transporters are required for ABA homeostasis under normal conditions, whereas the suggested conjugated-ABA-sink mechanism is restricted in abiotic stress environments, enabling rapid ABA responses at distinct target sites (Fig. 6j).

Our model raises several questions. It is unknown how free ABA, which is not taken into the shoot cortex or mesophyll cells, finds its way to the guard cells, which requires short-distance transport, and lateral-root formation sites, which requires long-distance transport. ABA might move through the apoplast, passively diffuse, or be actively translocated over the plasma membrane. Several ABA transporters have been identified so far. In *Arabidopsis* ABCG40 is the main ABA importer into guard cells ^15, 45^, and in tomato AIT1.1 was recently shown to play a role in ABA import into guard cells in a DELLA-dependent manner ^62^. In addition, NPF4.6, ABCG25 and DTX50 regulate vasculature ABA import and export ^25^. We speculate that it is unlikely that free ABA can travel from shoot to root by simple diffusion and that there are transporters that remain to be identified that mediate shoot-to-root ABA transport. These transporters are expected to load ABA into the shoot phloem and unload ABA in the root to regulate lateral root development. Our proposed model suggests that ABCG17 and ABCG18 mediate the rate-limiting step for ABA accumulation in shoots, thus allowing ABA translocation to roots while ABA-GE accumulates within the shoot cortex and mesophyll cells (Fig. 6j). However, it is currently unclear whether ABA-GE itself can travel from root to shoot and *vice versa ^30^* and further tools are required to characterize such mechanisms.

Previous studies have shown that ABA translocation from root to shoot via the xylem is required for stomatal closure, which restricts transpiration and water loss under drought stress (*12-16*). Little is known, however, about shoot-to-root transport via the phloem. McAdam *et al.* presented evidence that most of the ABA found in the tomato root is synthesized in the shoot ^31^. It is therefore possible that ABCG17 and ABCG18 are the mechanism regulating this process in *Arabidopsis*. Further studies in tomato and other species are needed to confirm whether the ABCG17- and ABCG18-mediated shoot-to-root ABA translocation mechanism is conserved. Ramzi *et al.* showed that the accumulation of ABA in roots after long-term water stress largely relies on shoot-to-root ABA transport ^63^, a process that is likely conserved in *Arabidopsis* ^17^, citrus ^64^ and other species. We show that *ABCG17* and *ABCG18* are downregulated under abiotic stress conditions, supporting the idea that their reduced activity promotes shoot-to-root ABA translocation under stress conditions.

The model proposed here suggests that ABCG17 and ABCG18 drive an inactive ABA-GE sink that restricts stomata activity and lateral root formation. However, we currently have little information on the cell-type or subcellular distributions of ABA and ABA-GE or of the associated ABA-glucosyltransferase localization. A detailed map of ABA-glucosyltransferase activity is required to plot ABA and ABA-GE movement in high resolution. Such a task remains challenging as this large family of enzymes is likely regulated at the protein level and profiling the expression pattern will not be sufficient. High-resolution fluorescent probes specific for ABA-glucosyltransferase that can be used in live plants would be an extremely useful tool in the field. There are some indications for ABA-GE accumulation in intracellular storage organelles (such as vacuoles) ^65^, but it remains unclear exactly how ABA and ABA-GE distribute within the mesophyll cells. We speculate that ABCG17 and ABCG18 activity leads to high levels of free ABA within the mesophyll and shoot cortex cells, which is then converted to ABA-GE by UDP-glycosyltransferases. Current models suggest that the ABA-GE is translocated to the endoplasmic reticulum and vacuole for storage ^45, 66^. Burla *et al.* suggested that two distinct membrane transport mechanisms are active in *Arabidopsis* mesophyll cells: a proton gradient-driven and an ATP-binding cassette transporter-mediated. They demonstrated that ABCC1 and ABCC2 are localized in tonoplasts and exhibit ABA-GE transport activity in a yeast heterologous expression system ^67, 68^. It is therefore possible that ABCC and ABCG transporters work in concert: ABCG17 and ABCG18 import free ABA into mesophyll cells, and ABCC1 and ABCC2 mediate storage of ABA-GE in vacuoles. In addition to the need to identify subcellular ABA-GE transporters, there is no direct evidence of ABA and ABA-GE distribution to different organelles. New biochemical and genetic tools are required to generate a comprehensive ABA profile at subcellular resolution.

## Methods

### Plant material and growth conditions

All *Arabidopsis thaliana* lines are in Col-0 background. For assays on plates, sterilized seeds were plated on 16 × 16 cm square petri dishes or 8.5 cm round petri dishes with growth media containing 0.5 × Murashige-Skoog (MS) medium (pH 5.95 - pH 6.1, 1% sucrose, 0.8% plant agar). The sterilized seeds were transferred to growth chambers (Percival, CU41L5) at 21 °C, 100 μEm^−2^ S^−1^ light intensity under long-day conditions (16 h light/8 h dark) after 3-day-stratification at 4 °C in the dark. For soil experiments, seedlings were transferred onto soil and grown in dedicated growth rooms under long-day conditions (16 h light/8 h dark) at 21 °C. The following ABA reporters were previously described: *pMAPKKK18:GUS, pMAPKKK18:LUC ^56^* and *pRAB18:GFP ^55^.*

### *Agrobacterium* transformation

GV3101 electro-competent *Agrobacterium tumefaciens* strain was incubated on ice with ~100 ng plasmid for 5 min and electroporated using a MicroPulser (Bio-Rad Laboratories; 2.2 kV, 5.9 ms). Immediately after electroporation, 600 μl LB medium was added, and sample was shaken for 2 h at 28 °C. The *Agrobacterium* was then plated on selective LB agar plates containing the relevant antibiotics for 2 days at 28 °C.

### *Arabidopsis* transformation

*Agrobacterium* vectors were validated by colony PCR and sequencing before growing in 100 ml LB medium containing 25 μg/ml gentamycin, 50 μg/ml rifampicin, and vector-specific antibiotic for ~36 hours at 28 °C. *Agrobacterium* was harvested by centrifugation for 10 min at 4000 rpm, supernatant was discarded, and the bacteria pellet was resuspended in 40 ml 5% sucrose + 0.05× MS + 0.02% Silwet L-77. *Arabidopsis* flowers were dipped into the bacterial solution for ~5 minutes. After dipping, plants were kept in the dark for ~18 hours and grown until seeds were harvested. T1 seeds were sown on 0.5 × MS media containing the appropriate antibiotics for selecting transgenic plants or on soil with 0.1% basta spraying on ~10-day-old plants.

### Genotyping

T-DNA insertion lines for single mutants ordered from Gabi Kat (https://www.gabi-kat.de) and The *Arabidopsis* Information Resource (https://www.arabidopsis.org/) are listed in Supplementary Table 1. Primers for the T-DNA insertion mutants genotyping were designed using the T-DNA Primer Design Tool powered by Genome Express Browser Server (GEBD) (http://signal.salk.edu/tdnaprimers.2.html). Homozygous mutants were characterized by PCR carried out with primers listed in Supplementary Table 2.

### Cloning

*ABCG17*, *ABCG18*, and *ABCG19* coding regions were amplified using Phusion High-fidelity Polymerase (New England Biolabs) from Col-0 cDNA using primers listed in Supplementary Table 3. Promoters of *ABCG17* and *ABCG18* were amplified from Col-0 DNA using Phusion High-fidelity Polymerase (New England Biolabs) using primers listed in Supplementary Table 3. Promoters of *ABCG17* and *ABCG18* are 1276 and 1580 bp long respectively, including 5′ UTR. *ABCG17*, *ABCG18*, and *ABCG19* coding regions and as well as *ABCG17* and *ABCG18* promoter fragments were cloned into pENTR/D-TOPO (Invitrogen K2400), verified by sequencing, and subsequently cloned into binary destination vectors using LR Gateway reaction (Invitrogen 11791). *p35S:YFP-ABCG17* and *p35S:YFP-ABCG18* were generated using the pH7WGY2 vector and were selected using spectinomycin in *E. coli* and hygromycin in plants. *p35S:ABCG17* and *p35S:ABCG18* were generated using pH2GW7 vector and selected using spectinomycin in *E. coli* and hygromycin in plants. *p35S:XVE:ABCG17* and *p35S:XVE:ABCG18* were generated using pMDC7 vector and selected using spectinomycin in *E. coli* and hygromycin in plants. *pABCG17:LUC* and *pABCG18:LUC* were generated using the pFlash vector and selected using spectinomycin in *E. coli* and gentamycin in plants. *pABCG17:NLS-YFP* and *pABCG18:NLS-YFP* were generated using R1-R2:NLS-YFP in pART27 vector and selected using spectinomycin in *E.coli* and kanamycin in plants. *pABCG17:GUS* and *pABCG18:GUS* were generated using pWGB3 vector, kanamycin and hygromycin in *E. coli* and hygromycin in plants. *p35S:amiRNA* lines were generated using the pH2GW7 vector, spectinomycin in *E. coli* and hygromycin in plants. We designed the amiRNA at the WMD3 website (http://wmd3.weigelworld.org/cgi-bin/webapp.cgi). The amiRNA sequences were synthesized by Syntezza Bioscience Ltd. and were cloned into the pUC57 vector with two borders for the Gateway system, and then we cloned the amiRNAs into the destination vector pH2GW7 using the Gateway system.

### Plant genetics

All transgenic lines were generated from the destination vectors transformed into Col-0. T1 seeds were collected and selected on 1/2 MS plates containing appropriate antibiotics or on soil with basta spraying. At least 10 independent lines for each construct were generated. Two representative homozygous lines were obtained for further characterization. To generate overexpression lines in the background of the ABA reporters, the homozygous *ABCG17* or *ABCG18* overexpression lines were crossed with the ABA reporter lines *pRAB18:GFP*, *pMAPKKK18:GUS* or *pMAPKKK18:LUC*. F1 heterozygous seeds for both constructs were used for analysis. As a control, the respective ABA reporters were crossed with Col-0 to generate F1 seeds. To generate double-knockdown line *mir17,18* in the background of the ABA reporters, the homozygous *mir17,18* line was crossed with *pMAPKKK18:GUS* to obtain F1 seeds. Heterozygous seeds were used for further analysis. As a control, the *pMAPKKK18:GUS* ABA reporter was crossed with Col-0 to generate F1 seeds.

### Phylogenetic tree

A phylogenetic tree of *Arabidopsis* ABCG family members, based on protein sequences, was constructed using sequences alignment by Clustal W (http://www.clustal.org/clustal2/e). The one neighbor-joining phylogenetic tree was constructed using MEGA7.0 (Molecular Evolutionary Genetics Analysis) software with 1000 bootstrap replications.

### Gas-exchange measurements

Gas exchange was measured using a LI-COR LI-6800 portable gas-exchange system. Plants were grown in short-day conditions (8 h light/16 h dark). Photosynthesis was induced under 150 μmol m^−2^ s^−1^ light with 400 μmol mol^−1^ CO_2_ surrounding the leaves. The amount of blue light was set to 10% of the photosynthetically active photon flux density. The flow rate was set to 200 μmol air s^−1^, and the leaf-to-air vapor pressure deficit was kept around 1kPa during the measurement. Leaf temperature was ~22 °C. Measurements were performed between 10:00 AM and 1:00 PM (three hours after the light was switched on.).

### Fluorescently labeled ABA synthesis

The synthesis and characterization of ABA-FL are described in the supplemental information (Supplementary Fig. S12-13).

### Fluorescently labeled ABA protoplast experiments

ABA-FL was dissolved in DMSO to 5 mM stock solution. Protoplasts isolation was carried as previously described ^69^ with the following adaptations: Protoplasts were treated with ABA-FL for 15 min at a final concentration of 5 μM ABA-FL, washed with 500 μl W5 solution (2 mM solution (pH 5.7) containing 154 mM NaCl, 125 mM CaCl2 and 5 mM KCl) three times, resuspended in 100 μl W5 solution, and mounted on slides for confocal microscopy.

### Stomatal aperture measurements

Stomatal aperture was determined using a rapid and almost permanent imprinting technique previously described (Geisler, *et al*, 2000). Leaves of similar size were cut from 25-day-old plants. The abaxial side of the leaves was attached to 0.5 ml silicone impression material (elite HD+, Zhermack Clinical). The leaves were removed after the silicone impression material dried. Subsequently, transparent nail varnish was placed on the epidermal side. The samples were placed on slides with cover-slips and imaged with a light microscope ^70^. The stomatal aperture size was quantified using Fiji software.

### RNA extraction, cDNA synthesis, and quantitative PCR

RNA was isolated using SV Total RNA Isolation System (Promega). For cDNA synthesis, a High-Capacity cDNA Reverse Transcription Kit (Applied Biosystems) was used according to manufacturer information. Fast SYBR Green Master Mix (Applied Biosystems) was used for quantitative PCR as described before ^71^. The primers used for quantitative PCR are listed in Supplementary Table 4.

### Thermal imaging

Thermal images were obtained using a FLIR T6xx series instrument. The camera was placed vertically ~50 cm above the plants. Leaf temperature was quantified using a customized region-of-interest tool implemented according to the manufacturer’s instructions.

### Histology, microscopy, and phenotypic characterization

Sections were made using the Vibratome technology as previously described ^72^. Sections were stained with 0.01% (g/ml) Fluorescent Brightener 28 in 1 × PBS for 20 minutes, washed with 1 × PBS three times (10 min each wash), and subsequently imaged with a Zeiss LSM780 confocal microscope. Plant shoot images were taken using a Nikon camera (WEM ED IF Aspherical Micro 1:1 Φ 62). For root growth assays, seedlings were grown on plates and scanned using HP Scanjet G3110. Leaf size and root length were measured using Fiji software (https://fiji.sc/).

### GUS staining and histology

Histochemical detection of GUS activity was carried out using 5-bromo-4-chloro-3-indolyl-β-D-glucuronide as a substrate as previously described ^50^. Samples were placed on slides with glass cover-slips and imaged using Zeiss binocular microscope. Histology of GUS stained samples was performed using protocol as described previously ^73^.

### Leaf water-loss assay

Leaves of 30-day-old plants were cut and weighed immediately to determine the fresh weight. Leaves were then exposed to room temperature air and weighed at 30 min, 60 min, 90 min, 120 min, 180 min, and 480 min.

### Bioluminescence LUC assays

*ABCG17* and *ABCG18* promoters were cloned into the pFlash vector using the Gateway system to generate *promoter:LUC* reporter constructs. Constructs were then transformed into Col-0 plants, and homozygous transgenic plants were treated with 10 mg/ml luciferin (Promega) and placed in the dark for 5 min. The LUC signal intensity was subsequently detected using the Biospace system (camera: IS1643N7056).

### Grafting assays

Six-day-old plants were used for grafting assay according to procedures described previously ^74^. Both of the two cotyledons were cut from seedlings, media used in the grafting assay contained 0.5% sucrose, and grafted plants grew vertically in short-day conditions.

### Radioactive ABA translocation and transport assays

ABA long-distance shoot-to-root translocation was measured using [^3^H]ABA. Shoots were treated with 2 μl of 1 μM [^3^H]ABA. After 24 h, roots were excised and placed into 3 ml of scintillation liquid (Opti-Fluor) for at least 24 h in darkness before monitoring in a liquid scintillation counter. Simultaneous [^14^C]ABA and [^3^H]IAA from tobacco (*N. benthamiana*) mesophyll protoplasts were analyzed as described ^75^. Tobacco mesophyll protoplasts were prepared 4 days after agrobacterium-mediated transfection of *p35S:YFP-ABCG17*, *p35S:YFP-ABCG18, p35S:YFP-ABCG19* or empty vector control. Relative export from protoplasts is calculated from exported radioactivity into the supernatant as follows: (radioactivity in the supernatant at time t = × min.) - (radioactivity in the supernatant at time t = 0)) * (100%)/ (radioactivity in the supernatant at t = 0 min.); presented are mean values from 8 ([^3^H]ABA) and 4 ([^14^C]IAA) independent transfections.

### Hormone quantification

Plant samples frozen in liquid nitrogen were grounded using motor and pestle. Around 200 mg of samples (shoot or root) was measured from ground powder and extracted with ice cold methanol/water/formic acid (15/4/1 v/v/v) added with isotope labelled internal standards (IS). Similar concentrations of internal standards (IS) of ABAs, auxins and gibberellins were added into samples and calibration standards. The samples were purified using Oasis MCX SPE cartridges (Waters) according to manufacturer’s protocol. The samples were injected on Acquity UPLC BEH C18 column (1.7 μm, 2.1×100 mm, Waters; with gradients of 0.1% acetic acid in water or acetonitrile), connected to Acquity UPLC H class system (Waters) coupled with UPLC-ESI-MS/MS triple quadrupole mass spectrometer for identification and quantification of hormones. The hormones were measured using MS detector, both in positive and negative mode, with two MRM transitions for each compound. External calibration curves were used for quantification, and calculated through Target Lynx (v4.1; Waters) software by comparing the ratios of MRM peak areas of analyte to that of internal standard. Heat map was generated using the BAR HeatMapper Plus tool (http://bar.utoronto.ca/ntools/cgi-bin/ntools_heatmapper_plus.cgi).

### Statistical analysis

Two-tailed Student’s t-test was performed whenever two groups were compared. Statistical significance was determined using Excel and Prism 7 (https://www.graphpad.com/scientific-software/prism/).

### Data availability

All the data supporting the findings of this study are available within the article and Supplementary Information files or from the corresponding author upon reasonable request.

## Acknowledgments

We thank Julian Schroeder (UCSD) and Sean Cutler (UCR) for sharing ABA reporter seeds and constructs, *pRAB18:GFP ^55^*, *pMAPKKK18:GUS*, and *pMAPKKK18:LUC ^55, 56^*. This work was supported by grants from the Israel Science Foundation (2378/19 to E.S.), the Human Frontier Science Program (HFSP—LIY000540/2020 to E.S.), the European Research Council (757683-RobustHormoneTrans to E.S., and 679189 GAtransport to R.W), the PBC postdoctoral fellowship (to Y.Z.) the ADAMA Center for Novel Delivery Systems in Crop Protection fellowship (to S.L.) and by the Swiss National Funds (31003A-165877/1 to M.G.). The authors declare no competing interests.

## References

1. Zhu J-K. Salt and drought stress signal transduction in plants. Annual review of plant biology 53, 247–273 (2002).

2. Cutler SR, Rodriguez PL, Finkelstein RR, Abrams SR. Abscisic acid: emergence of a core signaling network. Annual review of plant biology 61, 651–679 (2010).

3. Zhu J-K. Abiotic stress signaling and responses in plants. Cell 167, 313–324 (2016).

4. Zeevaart J, Creelman R. Metabolism and physiology of abscisic acid. Annual review of plant physiology and plant molecular biology 39, 439–473 (1988).

5. Duan L, et al. Endodermal ABA signaling promotes lateral root quiescence during salt stress in Arabidopsis seedlings. The Plant Cell, tpc. 112.107227 (2013).

6. Robbins NE, Trontin C, Duan L, Dinneny JR. Beyond the barrier: communication in the root through the endodermis. Plant Physiology, pp. 114.244871 (2014).

7. Chater CC, Oliver J, Casson S, Gray JE. Putting the brakes on: abscisic acid as a central environmental regulator of stomatal development. New Phytologist 202, 376–391 (2014).

8. Goldbach H, Goldbach E. Abscisic acid translocation and influence of water stress on grain abscisic acid content. Journal of Experimental Botany 28, 1342–1350 (1977).

9. Setter TL, Brun WA, Brenner ML. Abscisic acid translocation and metabolism in soybeans following depodding and petiole girdling treatments. Plant Physiology 67, 774–779 (1981).

10. Setter TL, Brun WA, Brenner ML. Effect of obstructed translocation on leaf abscisic acid, and associated stomatal closure and photosynthesis decline. Plant Physiology 65, 1111–1115 (1980).

11. Milthorpe F, Moorby J. Vascular transport and its significance in plant growth. Annual Review of Plant Physiology 20, 117–138 (1969).

12. Wilkinson S, Davies WJ. ABA-based chemical signalling: the co-ordination of responses to stress in plants. Plant, cell & environment 25, 195–210 (2002).

13. Sauter A, Davies WJ, Hartung W. The long-distance abscisic acid signal in the droughted plant: the fate of the hormone on its way from root to shoot. Journal of Experimental Botany 52, 1991–1997 (2001).

14. Zhang H, Zhu H, Pan Y, Yu Y, Luan S, Li L. A DTX/MATE-type transporter facilitates abscisic acid efflux and modulates ABA sensitivity and drought tolerance in Arabidopsis. Molecular plant 7, 1522–1532 (2014).

15. Kuromori T, Seo M, Shinozaki K. ABA Transport and Plant Water Stress Responses. Trends in plant science, (2018).

16. Ikegami K, Okamoto M, Seo M, Koshiba T. Activation of abscisic acid biosynthesis in the leaves of Arabidopsis thaliana in response to water deficit. Journal of plant research 122, 235 (2009).

17. Christmann A, Weiler EW, Steudle E, Grill E. A hydraulic signal in root-to-shoot signalling of water shortage. The Plant Journal 52, 167–174 (2007).

18. Holbrook NM, Shashidhar V, James RA, Munns R. Stomatal control in tomato with ABA-deficient roots: response of grafted plants to soil drying. Journal of Experimental Botany 53, 1503–1514 (2002).

19. Christmann A, Hoffmann T, Teplova I, Grill E, Müller A. Generation of active pools of abscisic acid revealed by in vivo imaging of water-stressed Arabidopsis. Plant physiology 137, 209–219 (2005).

20. Bauer H, et al. The stomatal response to reduced relative humidity requires guard cell-autonomous ABA synthesis. Current Biology 23, 53–57 (2013).

21. Koiwai H, Nakaminami K, Seo M, Mitsuhashi W, Toyomasu T, Koshiba T. Tissue-specific localization of an abscisic acid biosynthetic enzyme, AAO3, in Arabidopsis. Plant Physiology 134, 1697–1707 (2004).

22. Tan BC, et al. Molecular characterization of the Arabidopsis 9-cis epoxycarotenoid dioxygenase gene family. The Plant Journal 35, 44–56 (2003).

23. Park J, Lee Y, Martinoia E, Geisler M. Plant hormone transporters: what we know and what we would like to know. BMC biology 15, 93 (2017).

24. Kang J, et al. Abscisic acid transporters cooperate to control seed germination. Nature communications 6, (2015).

25. Lacombe B, Achard P. Long-distance transport of phytohormones through the plant vascular system. Current Opinion in Plant Biology 34, 1–8 (2016).

26. Boursiac Y, Leran S, Corratge-Faillie C, Gojon A, Krouk G, Lacombe B. ABA transport and transporters. Trends Plant Sci 18, 325–333 (2013).

27. Slovik S, Daeter W, Hartung W. Compartmental redistribution and long-distance transport of abscisic acid (ABA) in plants as influenced by environmental changes in the rhizosphere—a biomathematical model. Journal of Experimental Botany 46, 881–894 (1995).

28. Wilkinson S, Davies WJ. Xylem sap pH increase: a drought signal received at the apoplastic face of the guard cell that involves the suppression of saturable abscisic acid uptake by the epidermal symplast. Plant physiology 113, 559–573 (1997).

29. Dodd I, Tan L, He J. Do increases in xylem sap pH and/or ABA concentration mediate stomatal closure following nitrate deprivation? Journal of Experimental Botany 54, 1281–1288 (2003).

30. Jiang F, Hartung W. Long-distance signalling of abscisic acid (ABA): the factors regulating the intensity of the ABA signal. Journal of Experimental Botany 59, 37–43 (2007).

31. McAdam SA, Brodribb TJ, Ross JJ. Shoot-derived abscisic acid promotes root growth. Plant, cell & environment 39, 652–659 (2016).

32. Merilo E, et al. Stomatal VPD response: there is more to the story than ABA. Plant physiology 176, 851–864 (2018).

33. Kuromori T, Sugimoto E, Shinozaki K. Inter-tissue signal transfer of abscisic acid from vascular cells to guard cells. Plant Physiology, pp. 114.235556 (2014).

34. Merilo E, et al. Abscisic acid transport and homeostasis in the context of stomatal regulation. Molecular plant 8, 1321–1333 (2015).

35. Lee KP, Piskurewicz U, Turečková V, Strnad M, Lopez-Molina L. A seed coat bedding assay shows that RGL2-dependent release of abscisic acid by the endosperm controls embryo growth in Arabidopsis dormant seeds. Proceedings of the National Academy of Sciences 107, 19108–19113 (2010).

36. Seo M, Koshiba T. Transport of ABA from the site of biosynthesis to the site of action. Journal of plant research 124, 501–507 (2011).

37. Kang J, et al. Plant ABC transporters. The Arabidopsis Book, e0153 (2011).

38. Do THT, Martinoia E, Lee Y. Functions of ABC transporters in plant growth and development. Current opinion in plant biology 41, 32–38 (2018).

39. Kang J, et al. PDR-type ABC transporter mediates cellular uptake of the phytohormone abscisic acid. Proc Natl Acad Sci U S A 107, 2355–2360 (2010).

40. Kuromori T, et al. ABC transporter AtABCG25 is involved in abscisic acid transport and responses. Proc Natl Acad Sci U S A 107, 2361–2366 (2010).

41. Cho M, Lee SH, Cho H-T. P-Glycoprotein4 Displays Auxin Efflux Transporter-Like Action in Arabidopsis Root Hair Cells and Tobacco Cells. Plant cell, (2007).

42. Pawela A, Banasiak J, Biała W, Martinoia E, Jasiński M. Mt ABCG 20 is an ABA exporter influencing root morphology and seed germination of Medicago truncatula. The Plant Journal 98, 511–523 (2019).

43. Seo M, et al. Regulation of hormone metabolism in Arabidopsis seeds: phytochrome regulation of abscisic acid metabolism and abscisic acid regulation of gibberellin metabolism. Plant J 48, 354–366 (2006).

44. Kanno Y, et al. Identification of an abscisic acid transporter by functional screening using the receptor complex as a sensor. Proc Natl Acad Sci U S A 109, 9653–9658 (2012).

45. Ma Y, Cao J, He J, Chen Q, Li X, Yang Y. Molecular mechanism for the regulation of ABA homeostasis during plant development and stress responses. International journal of molecular sciences 19, 3643 (2018).

46. Nambara E, Marion-Poll A. Abscisic acid biosynthesis and catabolism. Annu Rev Plant Biol 56, 165–185 (2005).

47. Liu Z, et al. UDP-glucosyltransferase71c5, a major glucosyltransferase, mediates abscisic acid homeostasis in Arabidopsis. Plant Physiology 167, 1659–1670 (2015).

48. Chen T-T, et al. The Arabidopsis UDP-glycosyltransferase75B1, conjugates abscisic acid and affects plant response to abiotic stresses. Plant Molecular Biology 102, 389–401 (2020).

49. Hauser F, et al. A genomic-scale artificial microRNA library as a tool to investigate the functionally redundant gene space in Arabidopsis. The Plant Cell Online 25, 2848–2863 (2013).

50. Zhang Y, et al. A transportome-scale amiRNA-based screen identifies redundant roles of Arabidopsis ABCB6 and ABCB20 in auxin transport. Nature communications 9, 4204 (2018).

51. Murchie EH, Lawson T. Chlorophyll fluorescence analysis: a guide to good practice and understanding some new applications. Journal of experimental botany 64, 3983–3998 (2013).

52. Liu L, et al. ATP binding cassette transporters ABCG1 and ABCG16 affect reproductive development via auxin signalling in Arabidopsis. The Plant Journal, (2020).

53. Yadav V, Molina I, Ranathunge K, Castillo IQ, Rothstein SJ, Reed JW. ABCG transporters are required for suberin and pollen wall extracellular barriers in Arabidopsis. The Plant Cell, tpc. 114.129049 (2014).

54. Shanmugarajah K, et al. ABCG1 contributes to suberin formation in Arabidopsis thaliana roots. Scientific reports 9, 1–12 (2019).

55. Kim T-H, et al. Chemical genetics reveals negative regulation of abscisic acid signaling by a plant immune response pathway. Current Biology 21, 990–997 (2011).

56. Okamoto M, et al. Activation of dimeric ABA receptors elicits guard cell closure, ABA-regulated gene expression, and drought tolerance. Proceedings of the National Academy of Sciences 110, 12132–12137 (2013).

57. Shani E, et al. Gibberellins accumulate in the elongating endodermal cells of Arabidopsis root. Proc Natl Acad Sci U S A 110, 4834–4839 (2013).

58. De Smet I, Signora L, Beeckman T, Inzé D, Foyer CH, Zhang H. An abscisic acid-sensitive checkpoint in lateral root development of Arabidopsis. The Plant Journal 33, 543–555 (2003).

59. Geng Y, et al. A spatio-temporal understanding of growth regulation during the salt stress response in Arabidopsis. The Plant Cell 25, 2132–2154 (2013).

60. Zhao Y, et al. The ABA receptor PYL8 promotes lateral root growth by enhancing MYB77-dependent transcription of auxin-responsive genes. Science signaling 7, ra53–ra53 (2014).

61. Bloch D, Puli MR, Mosquna A, Yalovsky S. Abiotic stress modulates root patterning via ABA-regulated microRNA expression in the endodermis initials. Development 146, dev177097 (2019).

62. Shohat H, Illouz-Eliaz N, Kanno Y, Seo M, Weiss D. The tomato DELLA protein PROCERA promotes ABA responses in guard cells by upregulating ABA transporter. bioRxiv, (2020).

63. Manzi M, Lado J, Rodrigo MJ, Zacarías L, Arbona V, Gómez-Cadenas A. Root ABA accumulation in long-term water-stressed plants is sustained by hormone transport from aerial organs. Plant and Cell Physiology 56, 2457–2466 (2015).

64. Manzi Fraga MJ, Pitarch-Bielsa M, Arbona V, Gómez Cadenas A. Leaf dehydration is needed to induce abscisic acid accumulation in roots of citrus plants. (2017).

65. Dietz KJ, Sauter A, Wichert K, Messdaghi D, Hartung W. Extracellular β-glucosidase activity in barley involved in the hydrolysis of ABA glucose conjugate in leaves. Journal of Experimental Botany 51, 937–944 (2000).

66. Schroeder JI, Nambara E. A quick release mechanism for abscisic acid. Cell 126, 1023–1025 (2006).

67. Jarzyniak KM, Jasiński M. Membrane transporters and drought resistance–a complex issue. Frontiers in plant science 5, 687 (2014).

68. Burla B, Pfrunder S, Nagy R, Francisco RM, Lee Y, Martinoia E. Vacuolar transport of abscisic acid glucosyl ester is mediated by ATP-binding cassette and proton-antiport mechanisms in Arabidopsis. Plant Physiology 163, 1446–1458 (2013).

69. Wu F-H, Shen S-C, Lee L-Y, Lee S-H, Chan M-T, Lin C-S. Tape-Arabidopsis Sandwich-a simpler Arabidopsis protoplast isolation method. Plant methods 5, 16 (2009).

70. Geisler M, Nadeau J, Sack FD. Oriented asymmetric divisions that generate the stomatal spacing pattern in Arabidopsis are disrupted by the too many mouths mutation. The Plant Cell 12, 2075–2086 (2000).

71. Tal I, et al. The Arabidopsis NPF3 protein is a GA transporter. Nature communications 7, (2016).

72. Mitra PP, Loqué D. Histochemical staining of Arabidopsis thaliana secondary cell wall elements. JoVE (Journal of Visualized Experiments), e51381 (2014).

73. De Smet I, Chaerle P, Vanneste S, De Rycke R, Inzé D, Beeckman T. An easy and versatile embedding method for transverse sections. Journal of microscopy 213, 76–80 (2004).

74. Marsch-Martínez N, Franken J, Gonzalez-Aguilera KL, de Folter S, Angenent G, Alvarez-Buylla ER. An efficient flat-surface collar-free grafting method for Arabidopsis thaliana seedlings. Plant methods 9, 1–9 (2013).

75. Henrichs S, et al. Regulation of ABCB1/PGP1-catalysed auxin transport by linker phosphorylation. EMBO J 31, 2965–2980 (2012).

